# Learning Invariant Graph Representations for Cox Survival Modeling under Distribution Shifts

**DOI:** 10.64898/2025.11.30.691365

**Authors:** Ka Ho Ng, Chengshang Lyu, Anna Jiang, Yinhu Li, Lingxi Chen

**Author notes:** These authors contributed equally to this work (Ka Ho Ng, Chengshang Lyu).

## Abstract

Survival prediction from high-dimensional biomedical data is frequently compromised by distribution shifts across multi-center cohorts, where models trained on specific populations often rely on spurious correlations that fail to generalize to new environments. While recent independence-driven reweighting techniques attempt to mitigate this, they typically treat patients as isolated instances, neglecting the intrinsic topological structures and biological pathways shared within patient populations. To address this limitation, we propose InvGraphCox (Invariant Graph Cox), a novel framework that integrates graph-structured representation learning with robust survival modeling. InvGraphCox constructs a *k*-nearest-neighbor patient graph to capture local manifold structures and employs a Variational Graph Autoencoder (VGAE) combined with a cohort-wise alignment mechanism to learn low-dimensional patient embeddings that are invariant to site-specific biases. We comprehensively evaluate the framework across three distinct experimental settings: the Curated Top-100 Gene Benchmark for stable biomarker identification, large-scale, high-dimensional transcriptomic datasets (Ovarian and Breast Cancer) for unsupervised representation learning, and clinical datasets (Breast and Lung Cancer) involving mixed-type covariates. Experimental results demonstrate that InvGraphCox consistently outperforms state-of-the-art baselines in terms of discrimination, calibration, and risk stratification, confirming its ability to extract robust, biologically meaningful representations in heterogeneous healthcare settings.

## Introduction

Survival analysis, which estimates the time until an event of interest occurs, is a cornerstone of modern biomedical research and clinical decision-making (1). From stratifying patient risk to guiding personalized treatment plans, the ability to accurately predict survival outcomes based on high-dimensional covariates, such as transcriptomics or clinical features, is critical (2). The Cox Proportional Hazards (Cox PH) model remains the gold standard in this field due to its interpretability and flexibility (3). However, a fundamental challenge persists in deploying these models in real-world healthcare settings: distribution shift (4).

In multi-center studies, training data (e.g., from a developed region or specific hospital) often differs distributionally from testing data (e.g., a different demographic or sequencing platform) (5–9). Standard survival models (3), including deep learning variants (10, 11), typically assume independent and identically distributed (i.i.d.) data. When this assumption is violated, models often latch onto “spurious correlations”, the unstable features that correlate with survival in the training set but fail to generalize to new cohorts.

Recent advancements, such as Stable Cox (12), have attempted to mitigate this by employing independence-driven sample reweighting to decorrelate features and isolate stable variables. While effective, these methods largely treat patients as isolated samples, potentially overlooking the rich, non-linear topological structures inherent in patient populations. Patients with similar molecular profiles often share biological pathways and prognostic trajectories, forming a manifold structure that linear reweighting may fail to capture. To address this limitation, we propose InvGraphCox (Invariant Graph Cox), a novel framework that integrates graph-structured representation learning with robust survival modeling under distribution shifts. Unlike methods that focus solely on feature selection (3) or reweighting (12), InvGraph-Cox explicitly models the patient-patient similarity structure. By constructing a *k*-nearest-neighbor (KNN) graph in the gene expression space, we capture the local topology of the patient population. We then employ a Variational Graph Autoencoder (VGAE) to learn low-dimensional, robust patient embeddings. Crucially, we implement a cohort-wise alignment mechanism to ensure these embeddings remain invariant to site-specific distribution shifts.

We comprehensively evaluate InvGraphCox across distinct experimental scenarios to test its robustness under varying data conditions. First, we employ the Curated Top-100 Gene Benchmark (covering HCC [5 cohorts, *n* = 958], Breast Cancer [5 cohorts, *n* = 1980], and Melanoma [5 cohorts, *n* = 332]) to benchmark our method against state-of-the-art feature selection baselines in identifying stable biomarkers from pre-filtered candidates. Second, to assess the model’s capacity for high-dimensional representation learning, we utilize large-scale, multi-cohort transcriptomic datasets for Ovarian Cancer (OV, 10 cohorts, *n* = 2942) and Breast Cancer (BRCA, 5 cohorts, *n* = 1443), where the feature space is not restricted to pre-selected genes. Finally, to demonstrate generalizability beyond omics, we evaluate InvGraphCox on clinical data for Breast Cancer (4 cohort, *n* = 858) and Lung Cancer (9 cohort, *n* = 862), challenging the model to handle mixed-type covariates under distribution shifts. Across these diverse scenarios, InvGraphCox consistently outperforms existing methods—including Stable Cox, Cox PH, and parametric AFT models—in terms of C-index, calibration, and risk stratification, while invariance diagnostics confirm its ability to learn biologically meaningful representations that are robust to cohort-specific biases.

## Results

### Overview of InvGraphCox

In Figure 1A, InvGraphCox is designed to handle either high-dimensional genomic expression data or tabular clinical data alongside with patient survival times. The central engine processes these inputs to produce distribution-robust outputs. Unlike traditional models, InvGraphCox generates a graph-based survival model resilient to shifts, patient latent space embeddings, patient risk scores, and interpretable biomarker risk scores.

**Fig. 1.**
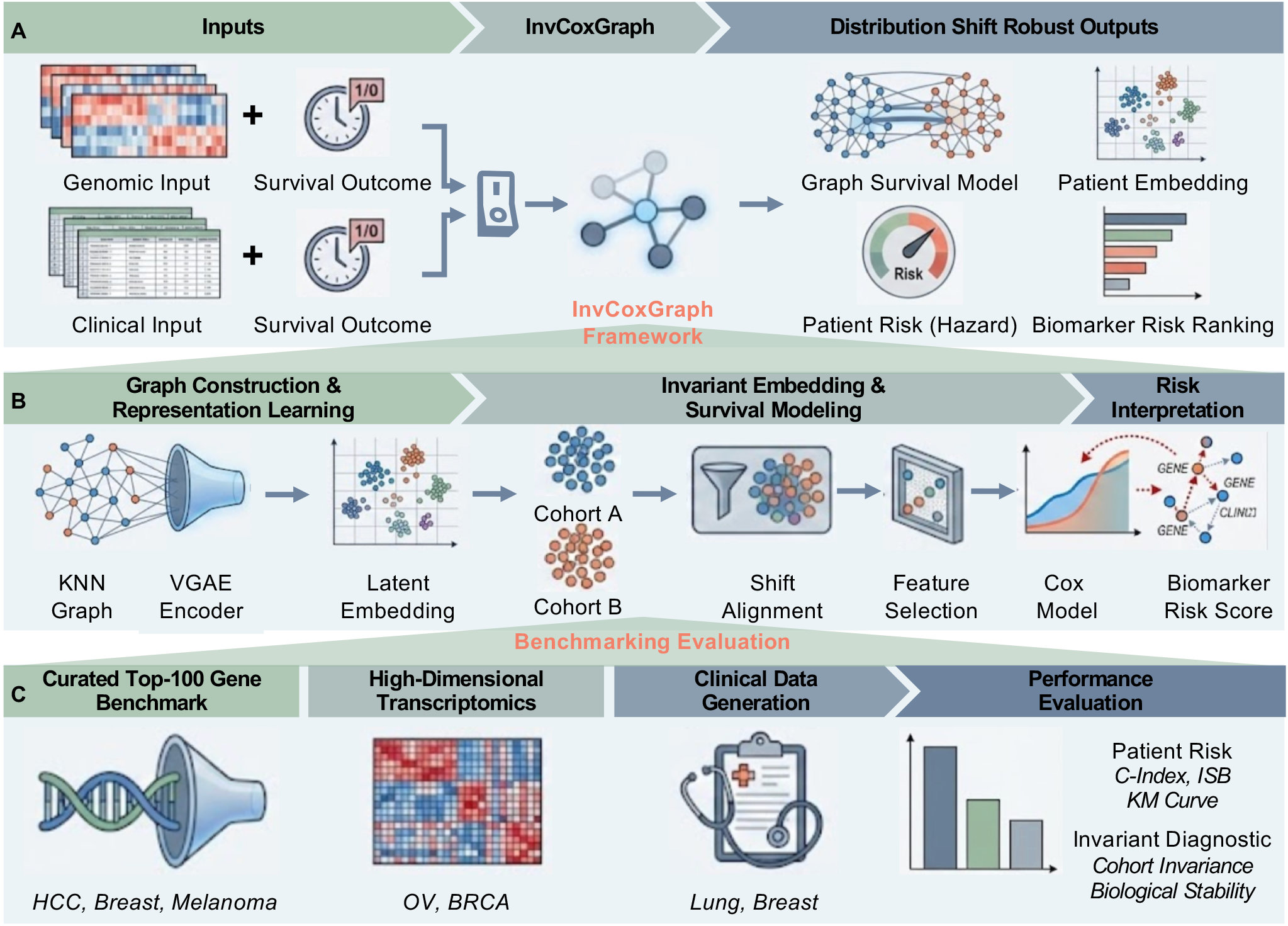
Graph overview of InvCoxGraph. **A** Model inputs and distribution robust outputs. **B** The InvGraphCox core algorithm. **C** Experimental benchmarking settings.

Figure 1B demonstrates that InvGraphCox employs a three-stage framework to learn robust, graph-aware patient representations and risk scores resilient to distribution shifts. The first stage is patient graph construction and representation learning. We first construct a patient-patient *k*-nearest-neighbors (KNN) graph within each cohort using Euclidean distances derived from gene expression or clinical data. To ensure structural comparability across cohorts despite covariate shifts, we fix the graph degree *k* and utilize a Gaussian kernel for edge weighting. A Variational Graph Autoencoder (VGAE) is then trained on the graph of the source cohort. By processing both node features *x* and neighborhood structures *N* (*v*), the VGAE learns low-dimensional latent embeddings *z* that encapsulate both feature information and topological context. The second stage is invariant embedding and survival modeling. To mitigate distribution shifts, we freeze the trained encoder and generate embeddings for all cohorts, applying cohort-wise residualization to align the mean embeddings of test cohorts with the training distribution. We then fit a penalized Cox proportional hazards model on these aligned embeddings. To enhance robustness and reduce variance, we perform a two-step feature selection: computing Wald *p*-values for all embedding dimensions and retaining only the top-*k* most significant dimensions to train the final Cox predictor. The third stage is gene-level risk attribution and interpretation. To bridge the gap between patient latent representations and clinical interpretability, we implement a back-mapping mechanism to assign risk scores to individual genes. We calculate the gradient of the predicted hazard ratio with respect to the input gene expression profile for each patient. By averaging these gradients across the aligned population, we derive a global gene risk score. This process transforms the “black box” embeddings into an interpretable ranking of high-risk genes, isolating stable biomarkers that maintain prognostic power across distributionally shifted cohorts.

Finally, we validate the framework across three distinct scenarios, as shown in Figure 1C: a curated benchmark using top-100 gene sets across HCC, Breast, and Melanoma datasets; evaluation on high-dimensional transcriptomics in Ovarian and BRCA cancer cohorts; and testing generalization on clinical data for Lung and Breast cancer. To rigorously emulate real-world distribution shifts in each scenario, we adopted a “leave-one-domain-out” strategy. Models were trained and tuned exclusively on a designated training cohort and a separate validation cohort. Evaluation was performed on disjoint, non-overlapping external testing cohorts. Importantly, no outcome information from the test cohorts was used during the training or hyperparameter tuning phases.

### Benchmarking Prognostic Performance on Curated Top-100 Gene Signatures

To rigorously evaluate the prognostic capability of InvGraphCox, we utilized the Stable Cox-preprocessed datasets (Top-100 genes and Overall Survival information) (12) spanning three distinct cancer types: Hepatocellular Carcinoma (HCC), Breast Cancer (BRCA), and Melanoma. These datasets pose significant challenges due to inter-cohort heterogeneity and varying sample sizes (detailed statistics provided in the Methods - Experiment Settings section). We benchmarked our model against the state-of-the-art Stable Cox (12), the classical Cox Proportional Hazards (Cox PH) model (3), and three parametric Accelerated Failure Time (AFT) models (Weibull, Log-logistic, and Log-normal).

We first assessed the model’s ability to predict patient risk scores by measuring the Concordance Index (C-index; detailed algorithm in the Methods - Concordance Index section) under stepwise Top-*K* gene thresholds (*K* ∈ {5, 10, 15, 20}). This multi-threshold design enables us to investigate the trade-off between feature quantity and the robustness of prognostic signals. As presented in Figure 2A (overall trends) and Figure 2B (cohort-specific details), InvGraphCox demonstrated superior predictive accuracy across most settings. Specifically, on the HCC and Breast Cancer datasets, our model achieved the highest mean C-index across all thresholds, yielding overall improvements of 4.43% and 1.96% over Stable Cox, respectively. Detailed cohort analysis reveals that InvGraphCox outperformed Stable Cox on all three HCC cohorts (increases of 5.71%, 2.49%, and 4.85%) and two out of three BRCA cohorts (8.43% and 1.84%). For the Melanoma datasets, a distinct performance trend was observed. While Stable Cox showed slightly better results at lower thresholds (Top 5 and 10), InvGraphCox surpassed it at higher thresholds (Top 15 and 20), achieving an 8.51% improvement at Top 20. Moreover, InvGraphCox exhibited a more stable performance across thresholds in the Melanoma, with average C-index variations of 0.0058 compared to Stable Cox’s 0.0172, suggesting enhanced robustness when incorporating a larger number of genes. This phenomenon highlights the impact of feature selection: suboptimal performance at lower thresholds likely stems from the presence of sparse biological signals in small gene sets. Conversely, while increasing the number of genes generally improves performance, surpassing an optimal point can introduce noise. Notably, Stable Cox exhibited a performance decline when the threshold reached 15, whereas InvGraphCox maintained robustness, effectively filtering noise through its graph-based constraints even as feature dimensionality increased.

**Fig. 2.**
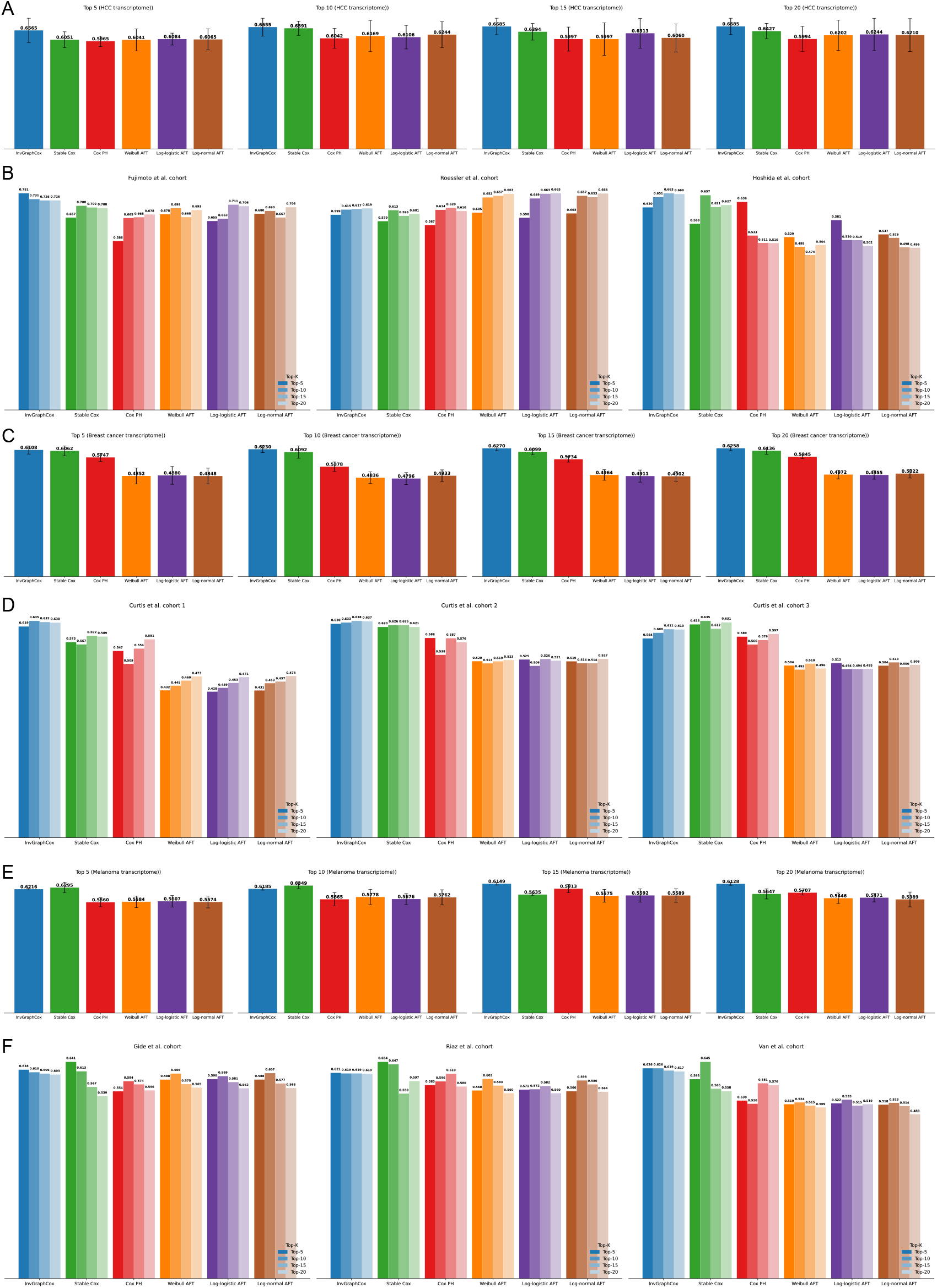
Performance Evaluation Based on C-index for All Cohorts of the Top-100 Gene Benchmark Data. **A** Mean C-index results for the top selected features across six methods trained on the TCGA-LIHC hepatocellular carcinoma transcriptome cohort (n=351). **B** C-index values for Fujimoto et al. (n=203), Roessler et al. (n=209), and Hoshida et al. (n=80) cohorts based on the top selected features (5, 10, 15, and 20) identified from the TCGA-LIHC hepatocellular carcinoma transcriptome data. **C** Mean C-index results for the top selected features across six methods trained on the training cohort of the breast cancer transcriptome (n=763). **D** C-index values for Curtis et al. cohort 1–3 (n=521, 288, 238) based on the top selected features identified from the training cohort of the breast cancer transcriptome data. **E** Mean C-index results for the top selected features across six methods trained on the Liu et al. cohort of the Melanoma transcriptome (n=120). **F** C-index values for Gide et al. cohort (n=91), the Riaz et al. cohort (n=54), and the Van et al. (n=41) cohorts based on the top selected features identified from the Liu et al. cohort of the Melanoma transcriptome data.

Building on the robust performance observed at higher feature dimensions, we fixed the feature selection to the Top-20 genes for a comprehensive survival analysis. Patients were stratified into high- and low-risk groups based on the median risk score (*β*^*T*^ *X*) to generate Kaplan-Meier (KM) curves (Figure 3A–F). InvGraphCox successfully achieved statistically significant risk stratification (*p<* 0.05, Log-rank test) across all cohorts in the HCC and BRCA datasets (Figure 3A, C). Notably, compared to Stable Cox, our model demonstrated superior prognostic capability—evidenced by lower *p*-values and higher Hazard Ratios (HRs)—across the HCC, BRCA, and Melanoma (Gide et al. and Van et al.) cohorts. While InvGraphCox consistently distinguished high-risk patients with reduced survival probabilities across all tested datasets, Stable Cox yielded significant stratification in only five out of nine cohorts (Figure 3B, D, F). This sharp contrast underscores the superior robustness of InvGraphCox in identifying high-risk patients across diverse data sources.

**Fig. 3.**
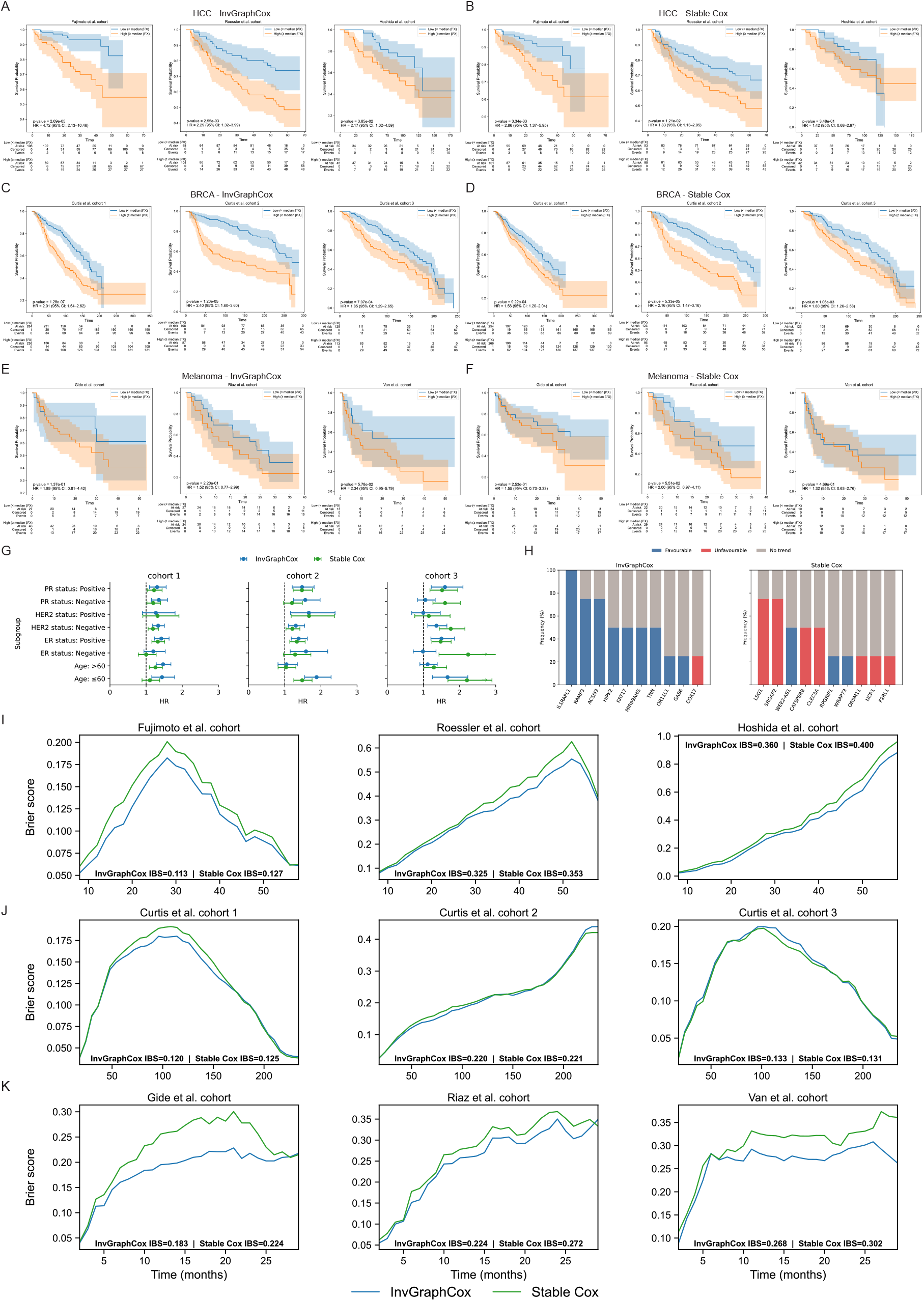
Survival and Performance Evaluation and Comparison of Prognostic Models for All Cohorts of the Top-100 Gene Benchmark Data. Kaplan-Meier (KM) survival plots (A–F) are shown for the indicated cancer cohorts. Subgroups are defined by the median of the risk score (*β*^*T*^ *X*) derived from the top 20 genes identified by the respective model (InvGraphCox or Stable Cox). **A** KM plots for three HCC test cohorts (InvGraphCox). **B** KM plots for three HCC test cohorts (Stable Cox). **C** KM plots for three BRCA test cohorts (InvGraphCox). **D** KM plots for three BRCA test cohorts (Stable Cox). **E** KM plots for three Melanoma cancer test cohorts (InvGraphCox). **F** KM plots for three Melanoma cancer test cohorts (Stable Cox). **G** Univariate Cox regression analysis of the hazard ratio (HR) across clinical subgroups for the three BRCA cohorts. The HR (box) and its 95% confidence interval (error bars) are calculated for refined subgroups. Refined subgroups are defined by an optimal cutoff value that ensures a minimum proportion of 30% of observations in each. **H** Prognostic consistency of individual genes across training and test cohorts of breast cancer. **I** Integrated Brier Score (IBS) comparison for three HCC test cohorts. IBS values for the InvGraphCox model are compared against the Stable Cox model over the survival follow-up time. Lower IBS indicates better model performance. **J** IBS comparison for three BRCA test cohorts. **K** IBS comparison for three Melanoma cancer test cohorts.

To evaluate the robustness of our model against specific clinical characteristics, we performed univariate Cox regression analyses across distinct clinical subgroups (including ER, PR, HER2 status, and Age), as shown in Figure 3G. InvGraphCox demonstrated superior discrimination, yielding higher Hazard Ratios (HRs) than Stable Cox in 16 of 24 subgroup-cohort comparisons. Notably, our model showed a particularly consistent prognostic signal in patients with positive PR and ER status across all cohorts. This suggests that InvGraphCox effectively resolves the heterogeneity within hormone receptor-positive (HR+) breast cancer, a population generally considered to have a favorable prognosis but containing a clinically critical high-risk subset (often associated with Luminal B-like profiles) (13, 14). By accurately identifying these high-risk patients despite their receptor positivity, InvGraphCox demonstrates strong clinical utility beyond standard markers.

We further assessed the biological consistency of the top 10 identified genes across the training and testing cohorts (Figure 3H). While both models maintained consistent directionality (no genes exhibited conflicting risk associations across cohorts), InvGraphCox achieved a significantly higher frequency of statistical significance. For instance, the gene *IL1RAPL1* showed a statistically detectable survival association across all cohorts (100% frequency) in our model, demonstrating robust generalization and reduced cohort-specific overfitting. Interestingly, we observed that InvGraphCox prioritizes “protective” biomarkers (associated with better prognosis, HR *<* 1), whereas Stable Cox tends to select “risky” biomarkers (HR *>* 1). This indicates a distinct difference in how the graph-based approach leverages biological signals compared to the re-weighting baseline.

To evaluate the accuracy of survival probability predictions over time, we calculated the Integrated Brier Score (IBS) (15) (Figure 3I–J), where a lower value indicates better calibration. InvGraphCox achieved consistently lower mean IBS values (0.266 ±0.109, 0.157 ±0.044, and 0.225 ±0.035 for HCC, BRCA, and Melanoma, respectively) compared to Stable Cox, particularly in the HCC and Melanoma cohorts (see blue vs. green lines in Figure 3I–J). These results confirm that InvGraphCox not only discriminates risk groups effectively but also provides more calibrated and accurate estimates of absolute survival probability.

Collectively, these results demonstrate that InvGraphCox achieves superior predictive accuracy and calibration across diverse cancer types, while also exhibiting remarkable robustness against cohort heterogeneity. By consistently outperforming Stable Cox in risk stratification and biological consistency, InvGraphCox establishes itself as a reliable and biologically interpretable tool for precision prognostic modeling in oncology.

To validate the robustness of InvGraphCox against distribution shifts, we conducted invariance diagnostics across all testing cohorts using four complementary metrics: Site-AUC (for both risk scores and embeddings), the Hilbert-Schmidt Independence Criterion (HSIC) (16), and the Silhouette score (17). Collectively, these metrics assess whether the model’s outputs retain spurious information about the data’s origin.

The risk-cohort site-AUC assesses the invariance of the final risk scores. An ideal site-AUC value of 0.5 suggests the risk score contains no usable information for predicting the data’s cohort of origin. As shown in Table 2, our InvGraphCox model consistently achieved site-AUC values very close to 0.5 (HCC: 0.482, BRCA: 0.481, Melanoma: 0.432), demonstrating that its risk predictions are largely site-invariant. The Stable Cox model showed slightly higher site-AUC values, particularly on the BRCA dataset (0.537), suggesting its risk scores may retain more site-specific information.

**Table 1.** Comparative Summary of Survival Prediction Models and Frameworks. Technical and theoretical comparison of the proposed InvGraphCox model against established semi-parametric (Stable Cox, Cox PH) and parametric Accelerated Failure Time (AFT) models (Weibull, Log-logistic, and Log-normal) used for prognostic prediction in transcriptomic survival data.

|  |  |  |  |  |  |  |
| --- | --- | --- | --- | --- | --- | --- |
| Tool | InvGraphCox | Stable Cox (12) | Cox PH (3) | Weibull AFT (7) | Log-logistic AFT (8) | Log-normal AFT (9) |
| Type | Hybrid: GNN<br>+ Semi-parametric Cox | Semi-parametric with<br>invariant feature learning | Semi-parametric | Parametric<br>(AFT/PH) | Parametric<br>(AFT) | Parametric<br>(AFT) |
| Core<br>Technique | Graph construction<br>+ VGAE embeddings + Cox regression | Sample reweighting<br>+ Weighted Cox regression | Partial likelihood | MLE | MLE | MLE |
| Distribution<br>Assumption | None | None | None | Weibull | Log-logistic | Log-normal |
| Hazard<br>Function<br>Shape | Follows Cox<br>PH assumption | Flexible,<br>proportional hazards | Monotonic,<br>proportional hazards | Monotonic | Non-monotonic | Bell-shaped,<br>non-monotonic |
| Special<br>Strength | Captures complex patient relationships;<br>adapts to cross-cohort shifts;<br>integrates omics & graph learning | Robust to<br>spurious correlations;<br>interpretable coefficients | Widely used,<br>interpretable | Dual PH/AFT<br>formulation | Captures mid-term<br>peak risk | Models early/late<br>survival differences |

**Table 2.** Comparative Diagnostics of Model Invariance in Multi-Cohort Transcriptomic Data. Diagnostic metrics quantify the independence between the models’ output (risk scores or latent representations) and the source cohort identity, comparing the proposed InvGraphCox model against the Stable Cox baseline across HCC, Breast Cancer, and Melanoma transcriptome cohorts.

| Metric (↓ better) | InvGraphCox | Stable Cox | Note |
| --- | --- | --- | --- |
| **Panel A: HCC Transcriptome** |  |  |  |
| Site-AUC (risk → cohort) | $0.482 \pm 0.036$<br>(DfR=0.036) | $0.462 \pm 0.015$<br>(DfR=0.090) | $\approx 0.5$ is better |
| Site-AUC ($z \rightarrow$ cohort) | $0.390 \pm 0.029$<br>(DfR=0.220) | — | $\approx 0.5$ is better |
| HSIC($z$, cohort) | $2.7 \times 10^{-7}$ | — | $0 \Leftrightarrow$ indep. |
| Silhouette($z \mid$ cohort) | $-0.110$ | — | $\approx 0$ or $< 0$ is good |
| **Panel B: Breast Cancer Transcriptome** |  |  |  |
| Site-AUC (risk → cohort) | $0.481 \pm 0.022$<br>(DfR=0.038) | $0.537 \pm 0.016$<br>(DfR=0.074) | $\approx 0.5$ is better |
| Site-AUC ($z \rightarrow$ cohort) | $0.385 \pm 0.015$<br>(DfR=0.230) | — | $\approx 0.5$ is better |
| HSIC($z$, cohort) | $4.0 \times 10^{-8}$ | — | $0 \Leftrightarrow$ indep. |
| Silhouette($z \mid$ cohort) | $-0.075$ | — | $\approx 0$ or $< 0$ is good |
| **Panel C: Melanoma Transcriptome** |  |  |  |
| Site-AUC (risk → cohort) | $0.432 \pm 0.075$<br>(DfR=0.136) | $0.416 \pm 0.029$<br>(DfR=0.168) | $\approx 0.5$ is better |
| Site-AUC ($z \rightarrow$ cohort) | $0.337 \pm 0.044$<br>(DfR=0.326) | — | $\approx 0.5$ is better |
| HSIC($z$, cohort) | $1.97 \times 10^{-6}$ | — | $0 \Leftrightarrow$ indep. |
| Silhouette($z \mid$ cohort) | $-0.052$ | — | $\approx 0$ or $< 0$ is good |

**Metric Definitions and Interpretation:** The metrics verify that learned features are **invariant** (independent) to the source cohort.
**Abbreviations:** ↓ better: Lower values indicate better performance; **Site-AUC**: Site-specific Area Under the Curve; **DfR**: Difference from Random; **HSIC**: Hilbert-Schmidt Independence Criterion; **z**: Latent representation/embedding vector from the GNN component.

The z-cohort site-AUC evaluates the invariance of the learned embeddings (z) before the final Cox model is applied. Our InvGraphCox model consistently yielded site-AUC values significantly below 0.5 (HCC: 0.390, BRCA: 0.385, Melanoma: 0.337). The associated low Distance-from-Random (DfR) values indicate that the embeddings are highly site-agnostic, effectively stripping away identifiable cohort information.

More critically, HSIC and Silhouette provide further evidence of the embedding’s site invariance. HSIC measures the dependence between embeddings (z) and cohort labels, with an ideal value of zero indicating independence. Across all datasets, the InvGraphCox model achieved extremely low HSIC values (2.7 ×10^−7^, 4.0 ×10^−8^, and 1.97 ×10^−6^ for HCC, BRCA, and Melanoma, respectively), confirming that the embedding representations are highly independent of the cohorts.

Corroborating this, the Silhouette score assesses how well the embeddings cluster by cohort. Ideal values are near 0 or negative, indicating inter-cohort mixing rather than distinct clustering. Our results show negative Silhouette scores for all datasets (HCC: −0.110, BRCA: −0.075, Melanoma: −0.052), confirming that the model’s embeddings do not form tight clusters based on the original cohort. This strong mixing indicates that site-specific biases have been effectively removed from the latent representation.

The diagnostic results demonstrate that our InvGraphCox model successfully learns a site-invariant representation. The consistent achievement of ideal values across all four metrics and three distinct transcriptome datasets confirms that both the model’s embeddings and final risk scores are minimally influenced by the data’s original site or cohort.

Beyond site invariance, generalizability requires that the learned prognostic signals reflect shared biological mechanisms rather than cohort-specific artifacts. We assessed this using three complementary diagnostics: Top-K Jaccard overlap, Genome-wide rank Spearman correlation (18), and a Top-N signed Spearman heatmap.

In the Top-K Jaccard overlap assessment, we selected the top 20 genes with the highest gene-to-risk correlation score. Then we calculated the Jaccard overlap for every pair of cohorts and compared the results to a random baseline. As shown in Figure 4A, the Jaccard overlap for the Breast cancer transcriptome was consistently high, with a mean overlap of 0.562, far exceeding the random baseline. The HCC and Melanoma datasets also showed overlaps (mean 0.167 and 0.150, respectively) that were above their corresponding random baselines. Notably, the highest Jaccard score of 0.212 was observed in the comparisons of test1 vs. test2 for HCC, as well as in the test1 vs. test3 and test2 vs. test3 comparisons in the Melanoma datasets. These results demonstrate that our model consistently leverages a shared set of important biological features across different cohorts, indicating a stable and reproducible signal.

**Fig. 4.**
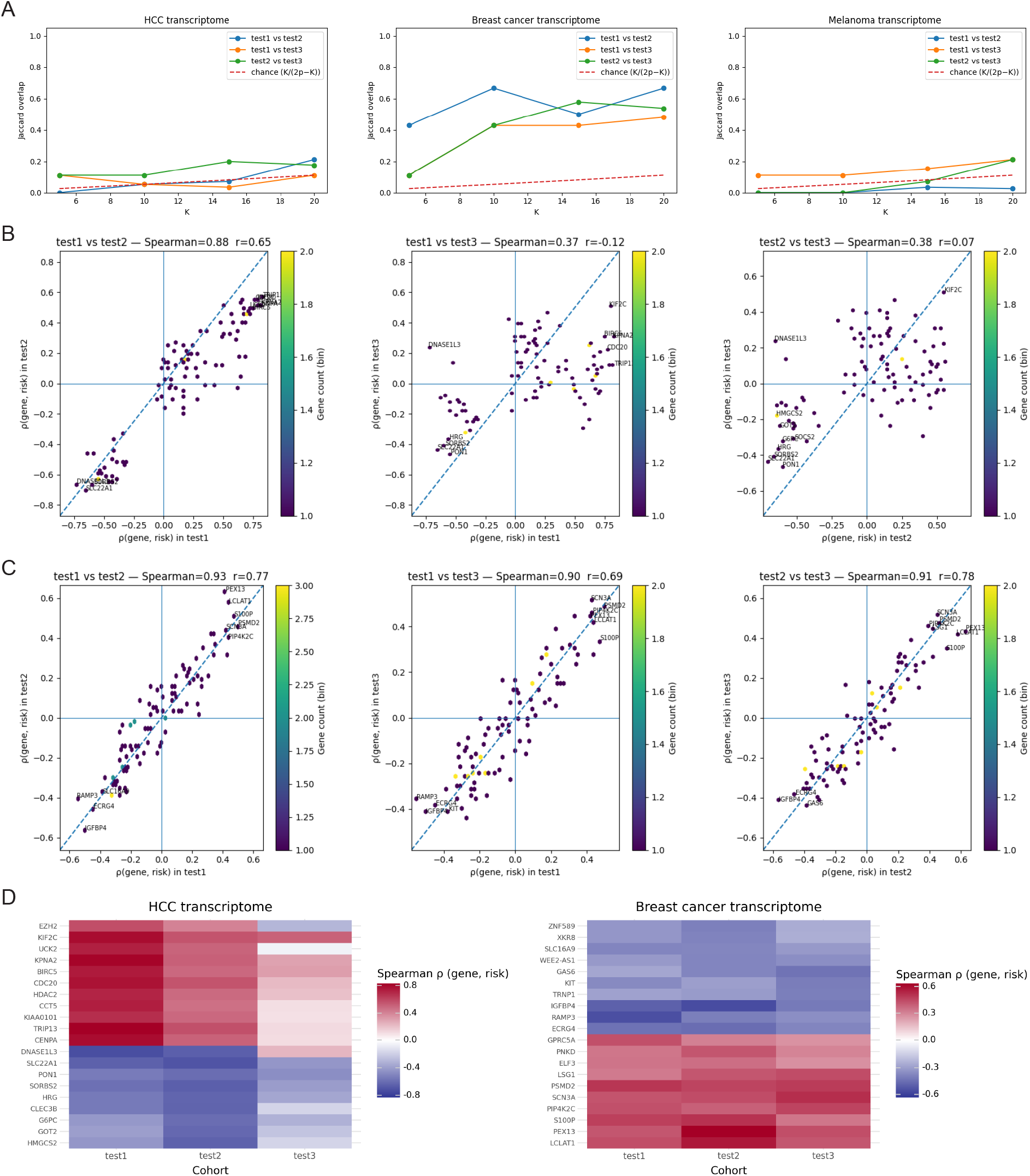
Stability and Consistency of Prognostic Gene Signatures in the Top-100 Gene Benchmark Data. The stability of the selected gene signature is evaluated across three independent test cohorts (test1, test2, test3) for the HCC, BRCA, and Melanoma datasets. **A** Jaccard Overlap of top-*K* ranked prognostic genes among pairs of the HCC, BRCA, and Melanoma test cohorts. The dashed line represents the expected overlap by chance. **B** Correlation of gene prognostic risk scores *p*(gene, risk) among HCC test cohorts. **C** Correlation of gene prognostic risk scores *p*(gene, risk) among BRCA test cohorts. **D** Consistency Heatmap of Gene Prognostic Effect (Spearman *ρ*) for the top-20 genes across the three HCC and BRCA test cohorts. Red indicates an unfavorable (high risk) association; Blue indicates a favorable (low risk) association.

Moving beyond the top features, we evaluated the global consistency of gene-to-risk associations using Genome-wide Rank Spearman Correlation (HCC and BRCA only; Melanoma was excluded due to limited sample size). The scatter plots in Figure 4B–C visualize these relationships. Each point represents a gene, plotted according to its correlation with risk in two different cohorts. A tight clustering of points along the 45-degree line (*y* = *x*) indicates a high level of agreement. We observed high mean genome-wide correlations for both HCC (*ρ* = 0.544) and BRCA (*ρ* = 0.912), with peak correlations reaching 0.879 and 0.926, respectively. This confirms that the global ordering of prognostic features remains robust across diverse data sources.

Moreover, the Signed Spearman Heatmap in Figure 4d illustrates the direction and magnitude of the gene-to-risk associations for the top-20 “meta-genes” in the HCC and Breast Cancer data. Both datasets exhibit remarkable consistency, with distinct blocks of genes maintaining a consistent sign and magnitude across all three cohorts. Specifically, the HCC dataset exhibits a sign consistency of 0.85 (17/20 genes) with low variance (0.080), whereas the BRCA dataset demonstrates perfect consistency (1.00) with negligible variance (0.004). This implies that InvGraphCox successfully captures the fundamental biological directionality of survival drivers.

Finally, we conducted an ablation study to compare the full InvGraphCox model (VGAE with SAGEConv) against three ablated variants: VGAE with GCNConv, GAE with SAGEConv, and the No GNN model (equivalent to a traditional penalized Cox model). This analysis was performed across three independent transcriptomic survival datasets (HCC, Breast Cancer, and Melanoma) to assess the robustness of each method across multiple external cohorts. Performance was evaluated using the Integrated Brier Score (IBS), site-level AUC, and C-index. The IBS results, shown in Figure 5A, demonstrate the superior calibration of InvGraphCox. Our model yields the lowest median Integrated Brier Score across all three datasets, indicating the most accurate calibration of survival probabilities over time. Additionally, the substantially lower variance in the IBS distribution for InvGraph-Cox compared to other baseline models suggests enhanced robustness to censoring and cohort shift. As shown in Figure 5B, the InvGraphCox model achieves the highest mean site-level AUC across the datasets. This result is interpreted as indicating a stronger ability to discriminate between high-risk and low-risk patients when the model is applied to independent testing cohorts. The C-index results, reported under the Top-*k* feature threshold, confirm the model’s predictive power. As highlighted in Figure 5C, InvGraphCox consistently achieves the highest C-index across all three transcriptomic datasets and throughout varying Top-*k* feature thresholds, further validating its strong prognostic performance.

**Fig. 5.**
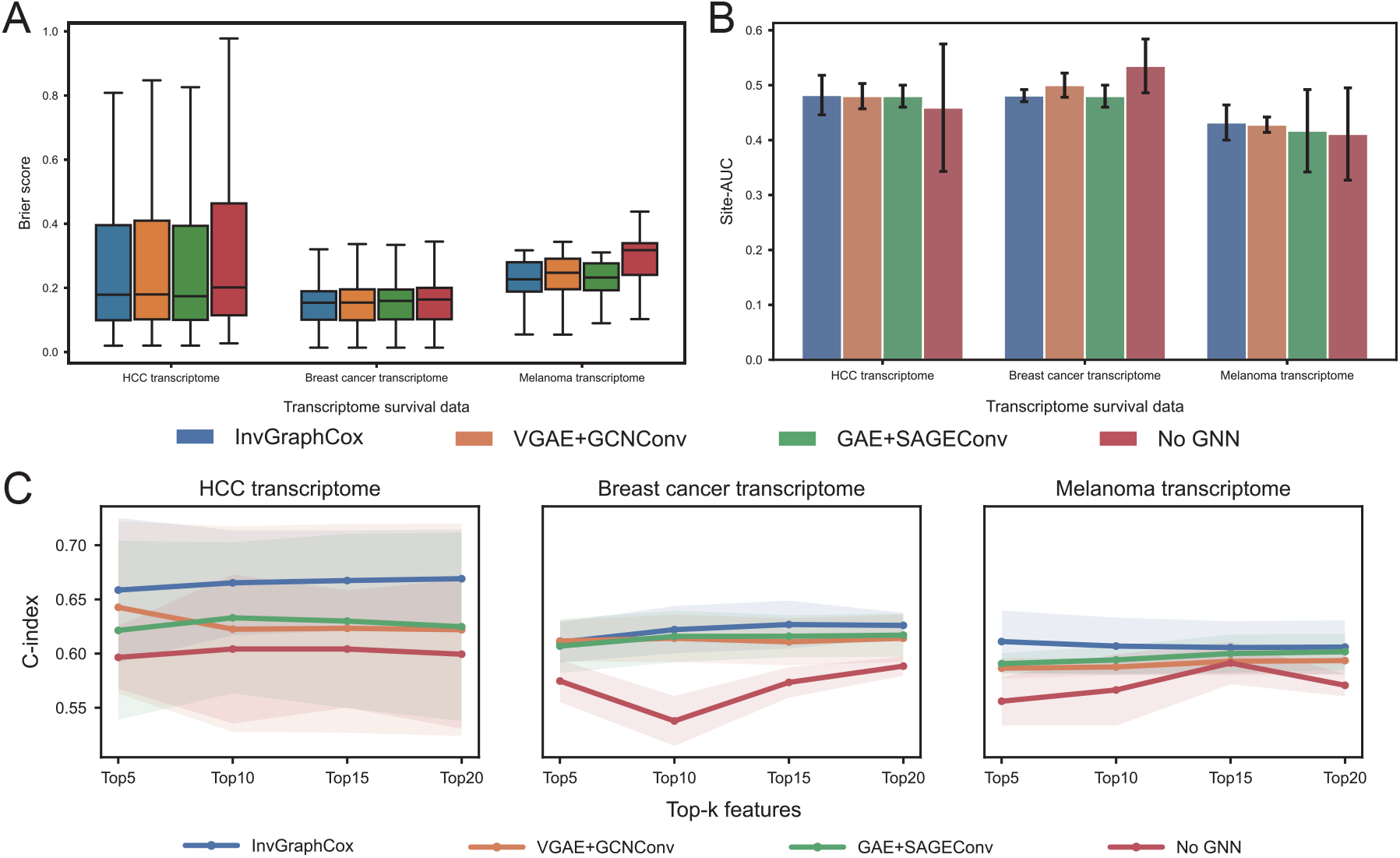
Ablation Study of the InvGraphCox Model Components in the Top-100 Gene Benchmark Data. Comparison of the prognostic performance of the full InvGraphCox model against three ablated versions: VGAE+GCNConv (using a standard GCN convolution), GAE+SAGEConv (using a Graph Autoencoder with SAGE convolution), and No GNN (removing the Graph Neural Network component entirely, equivalent to a traditional penalized Cox model) across three cancer types. **A** Integrated Brier Score (IBS) Comparison. Box plots show the distribution of the Integrated Brier Score (IBS) for the four model variants. A lower Brier Score indicates better overall prediction accuracy. **B** Site-AUC Performance. Bar plots comparing the mean Site-AUC value for each model variant across the three cancer types. Site-AUC measures the model’s ability to predict survival outcomes when applied to test cohorts, independent of the training cohort, indicating its cross-cohort generalization ability. **C** C-index versus Top-k Features. Line plots showing the Concordance Index (C-index) for the four model variants as a function of the number of top-ranked features (Top5, Top10, Top15, Top20) used in the risk score calculation. The shaded region indicates the standard deviation or confidence interval. A higher C-index indicates better discrimination ability.

### Robust Generalization to High Dimensional Transcriptomics Cohorts under Distribution Shifts

To rigorously evaluate the generalization capability of InvGraphCox under significant distribution shifts, we extended our validation to two large-scale multi-cohort transcriptome benchmarks: Ovarian Cancer (OV) and Breast Cancer (BRCA). For model training, we utilized the TCGA-OV dataset (*n* = 562) and Yau et al.’s breast cancer study (19) (*n* = 682). To assess robustness against cross-cohort heterogeneity, we evaluated the trained models on nine independent external cohorts for the OV dataset and three independent external cohorts for the BRCA dataset (details in the Methods - Experiment Settings). We benchmarked InvGraphCox against five baseline methods: Stable Cox, Cox PH, and three parametric AFT models (Weibull, Log-logistic, Log-normal). Performance was assessed using two complementary metrics: Concordance Index (C-index) and Kaplan-Meier (KM) Analysis.

For the Ovarian Cancer dataset, InvGraphCox demonstrated superior robustness and predictive accuracy across all tested feature thresholds (Top 5, 10, 15, and 20). As shown in Figure 6A, our model consistently achieved the highest C-index across the nine testing cohorts, yielding a mean C-index of 0.591 and a mean variation of 0.0283. This performance represents a substantial advantage over the state-of-the-art Stable Cox model (mean C-index 0.553, mean variation 0.0392) and the conventional Cox PH baseline (mean C-index 0.532, mean variation 0.0502). The traditional parametric AFT models showed comparatively weaker generalization, with C-index values ranging from 0.514 to 0.521. We further validated these findings using Kaplan-Meier survival analysis. Figure 6B illustrates the survival curves stratified by InvGraphCox for representative OV cohorts according to various platforms (e.g., GPL570, GPL7264, GPL7264). InvGraphCox successfully identified high-risk patient subgroups with significantly worse survival outcomes (e.g., GPL570: HR=1.45, *p<* 0.001; GPL2986: HR=1.96, *p<* 0.05; GPL7264: HR=1.90, *p<* 0.001). In contrast, the stratification produced by Stable Cox (Figure 6C) showed significant results in three out of nine cohorts. Although Stable Cox achieved significant stratification in some cohorts (e.g., GPL2986: HR=1.93, *p<* 0.05; GPL7264: HR=1.32, *p<* 0.2), its hazard ratios were consistently lower than those of InvGraphCox. These results underscore the superior prognostic capability of InvGraphCox in identifying high-risk patients across diverse challenging ovarian cancer cohorts.

**Fig. 6.**
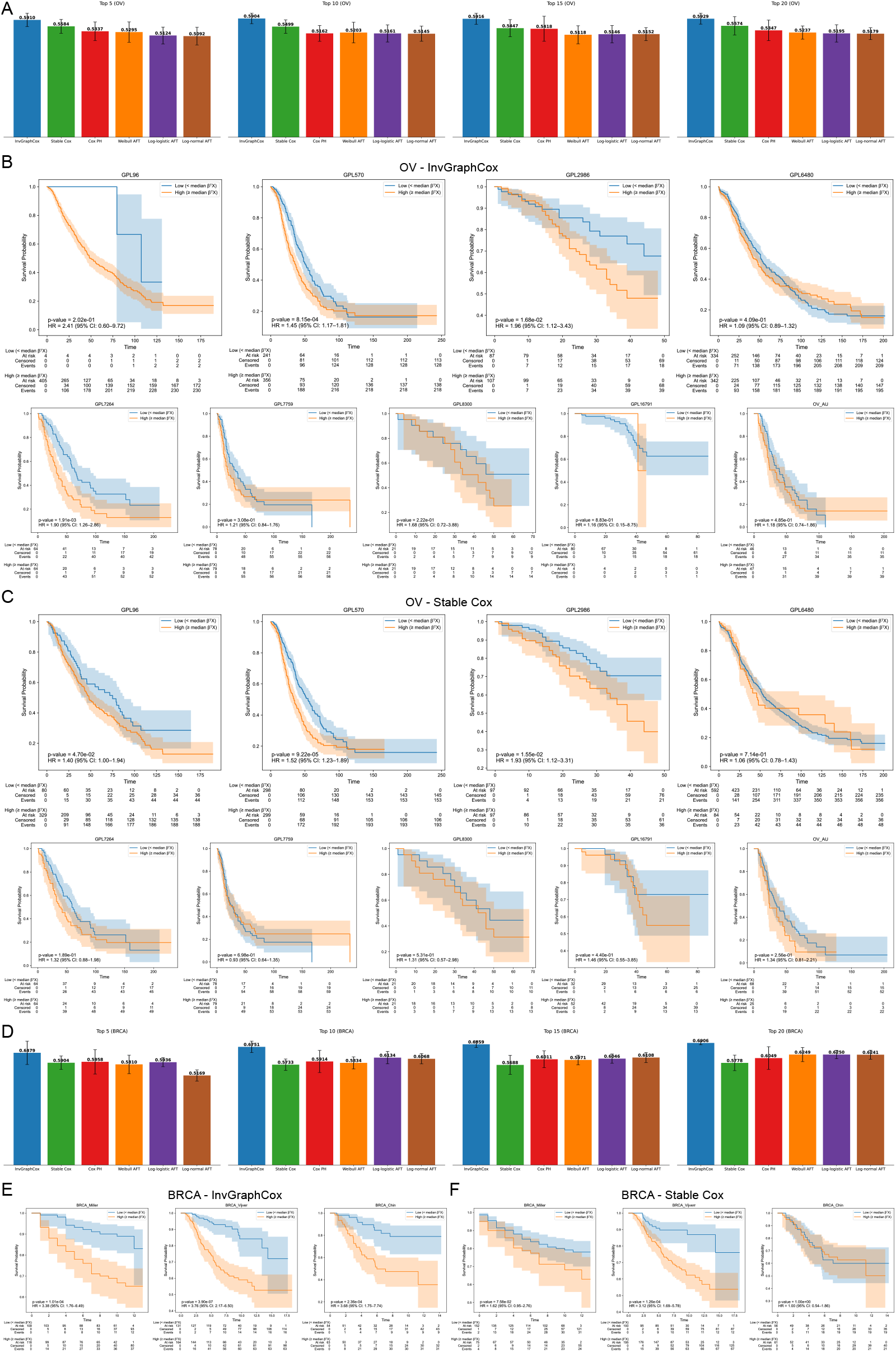
External Validation and Performance Comparison on Ovarian and Breast Cancer Cohorts. The prognostic performance of the InvGraphCox model is compared with the Stable Cox model using the High-Dimensional Transcriptomics Benchmark Data for Ovarian Cancer (OV) and Breast Cancer (BRCA). **A** C-index Comparison versus Top-*k* Features (Ovarian Cancer). Bar plots showing the C-index performance of the compared models as a function of the number of top-ranked features (Top 5 to Top 20) used in the risk score calculation for OV. A higher C-index indicates better discrimination. **B** Kaplan-Meier Curves (Ovarian Cancer, InvGraphCox). KM plots showing survival stratification results generated by the InvGraphCox model across various OV independent cohorts. **C** KM Curves (Ovarian Cancer, Stable Cox). KM plots showing survival stratification results generated by the Stable Cox model across various OV independent cohorts. **D** C-index Comparison versus Top-*k* Features (Breast Cancer). Bar plots showing the C-index performance of the compared models as a function of the number of top-ranked features (Top 5 to Top 20) used in the risk score calculation for BRCA. **E** KM Curves (Breast Cancer, InvGraphCox). KM plots showing survival stratification results generated by the InvGraphCox model across various BRCA independent cohorts. **F** KM Curves (Breast Cancer, Stable Cox). KM plots showing survival stratification results generated by the Stable Cox model across various BRCA independent cohorts.

For the Breast Cancer dataset, we conducted a granular analysis of performance improvements. The C-index comparisons across Top-*k* thresholds are summarized in Figure 6D. On the overall evaluation (mean across Test cohorts 1–3), InvGraphCox emerged as the top-performing method at every threshold, outperforming Stable Cox by margins of 9.72% (Top 5), 17.75% (Top 10), 20.59% (Top 15), and 19.53% (Top 20). Cohort-specific analysis reveals that InvGraphCox is consistently dominant on Test 2 (improvements of +11.00% to +15.53%) and particularly effective on Test 3, achieving dramatic gains (+25.48% to +40.26%). On Test 1, while InvGraphCox slightly underperformed Stable Cox at the lowest threshold (Top 5, −11.44%), it surpassed the baseline at higher thresholds (Top 10 to 20, improving by +14.10% to +15.04%). These trends suggest that while extremely small gene sets (Top 5) may omit informative signals necessary for generalizing to specific cohorts, such as Test 1, moderate thresholds (Top 10 to 20) allow InvGraphCox to incorporate richer biological signals, thereby maximizing generalization. The Kaplan-Meier plots further corroborate these quantitative gains. Figure 6E shows that InvGraphCox achieves clear and statistically significant separation between risk groups across the BRCA test cohorts. Conversely, Stable Cox (Figure 6F) exhibits weaker stratification, with survival curves that often overlap or lack statistical significance, and only one cohort shows significant divergence. Overall, InvGraphCox demonstrates robust gains over Stable Cox and other baselines, establishing itself as a reliable tool for cross-cohort breast cancer prognosis.

### Generalization to Survival Prediction with Clinical Mixed-Type Covariates

To assess the versatility of InvGraphCox beyond high-dimensional omics data, we evaluated its performance on multi-cohort clinical datasets characterized by mixed-type covariates (e.g., demographics, disease staging, treatment codes, and laboratory values). We focused on two major cancer types: Lung Cancer (endpoints: Overall Survival [OS], Disease-Free Survival [DFS]) and Breast Cancer (endpoints: OS, Recurrence-Free Survival [RFS]). Patients were partitioned into a single training cohort and distinct external test cohorts per endpoint to rigorously test out-of-distribution generalization.

We first compared the discrimination performance (C-index) across clinical subpopulations. As shown in Figure 7A for Lung Cancer, InvGraphCox achieved the best overall performance for OS, surpassing Stable Cox (0.690 vs. 0.661, an improvement of +4.36%). For DFS, Stable Cox performed slightly better (0.685 vs. 0.666). For Breast Cancer (Figure 7B), the two models were tied for OS (mean C-index ≈0.686), while InvGraphCox demonstrated a clear lead in RFS (0.685 vs. 0.666, +2.84%). In summary, InvGraphCox matches or exceeds the performance of the state-of-the-art Stable Cox in three out of the four clinical scenarios (Lung OS, Breast OS, Breast RFS).

**Fig. 7.**
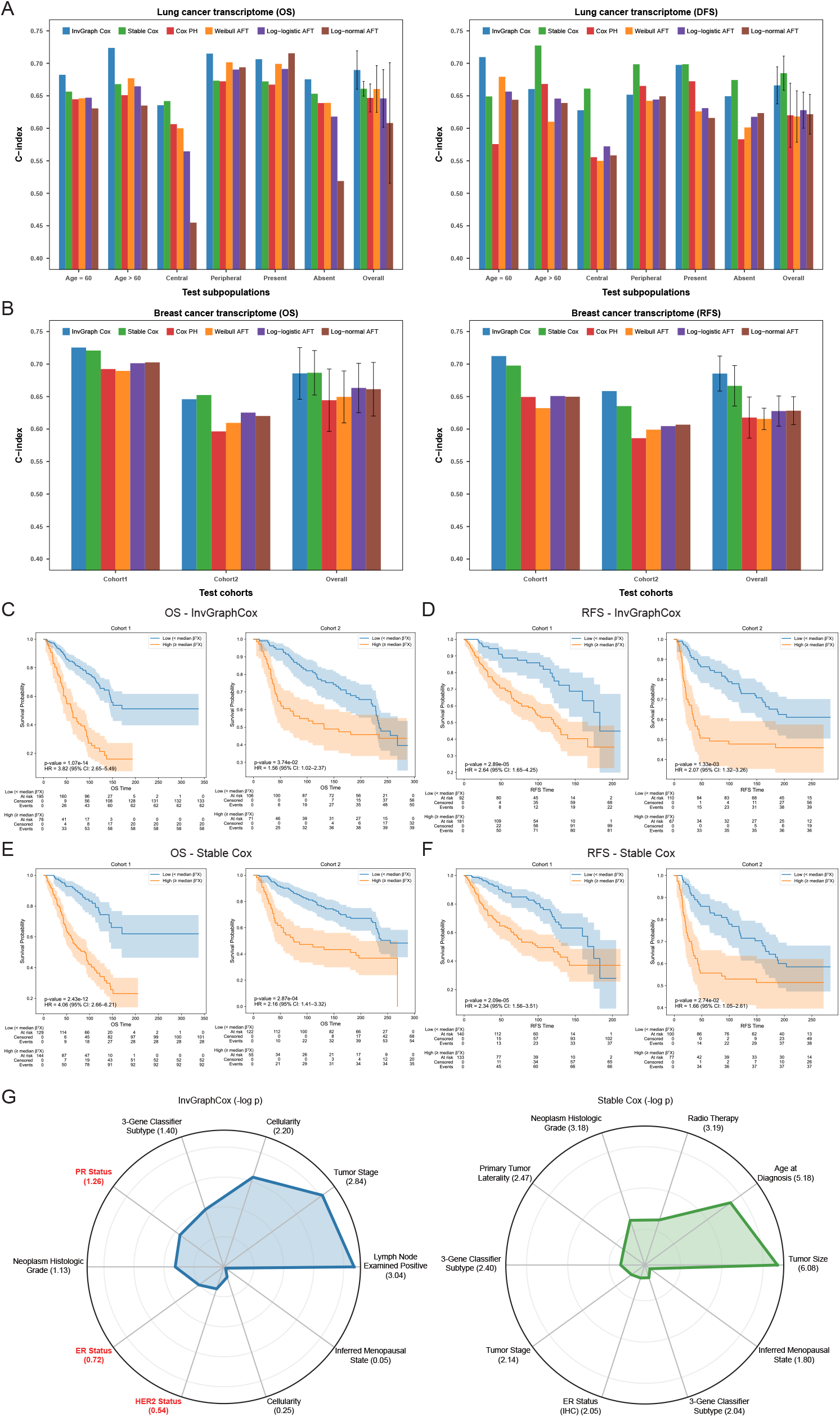
Comparative Performance of InvGraphCox in Clinical Survival Benchmark Data. **A** C-index Comparison in Lung Cancer Subpopulations (OS/DFS). Bar plots showing the C-index performance of the compared models for Overall Survival (OS) (left) and Disease-Free Survival (DFS) (right) across six predefined clinical subpopulations. A higher C-index indicates better discrimination. **B** C-index Comparison in Breast Cancer Cohorts (OS/RFS). Bar plots showing the C-index performance of compared methods, trained on the training cohort (*n* = 394), when evaluated on two independent test cohorts (Cohort 1, *n* = 273; Cohort 2, *n* = 177) for OS and Recurrence-Free Survival (RFS). The ‘Overall’ bar represents the mean C-index across all test cohorts. Error bars indicate the mean ± standard deviation (s.d.) of the results across the test cohorts, demonstrating the stable performance of the methods. **C** KM survival curves for OS in BRCA cohort 1&2., stratified by the InvGraphCox risk score (*β*^*T*^ *X*). **D** KM survival curves for RFS in BRCA cohort 1&2, stratified by the InvGraphCox risk score. **E** KM survival curves for OS in BRCA cohort 1&2, stratified by the Stable Cox risk score. **F** KM survival curves for RFS in BRCA cohort 1&2, stratified by the Stable Cox risk score. **G** Importance of Clinical Variables (Radar Plots). Radar plots showing the negative logarithm of the *p*-value (− log *p*) for each clinical variable derived from InvGraphCox (left, blue) and Stable Cox (right, green). Variables with larger radii (higher − log *p* values) are considered more prognostically significant.

The performance margins on clinical data are generally narrower and more cohort-dependent compared to our transcriptomic experiments. A plausible reason is that clinical data often contain many sparse one-hot covariates, such as staging and treatments. Even after z-scaling, these dimensions can still dominate Euclidean distances, causing KNN neighborhoods to reflect shared codes rather than prognostic similarity. That weakens graph homophily and reduces the benefit of message passing for a VGAE encoder. Moreover, because the graph kernel width (*σ*) is calibrated on training distances, shifts in the mix of binary vs continuous features across cohorts can misalign edge scales at test time. Together, these factors compromise the out-of-distribution advantage of graph-based representation on clinical covariates, explaining the smaller or mixed gains observed here compared to Stable Cox.

Following the procedure for an omics study, the Kaplan-Meier plots for high-risk and low-risk subgroups, as determined by the InvGraphCox model and the Stable Cox model for the breast cancer dataset, are presented in Figure 7C–F. Both InvGraphCox and Stable Cox significantly differentiated the two subgroups in all cohorts of the cancer dataset based on overall survival (OS) and recurrence-free survival (RFS) times. However, InvGraphCox demonstrated higher hazard ratios (HR) across three cohorts. This suggests that InvGraphCox is more effective at stratifying patients into distinct risk groups, which is crucial for personalized treatment planning and enhancing patient outcomes. The elevated HR scores suggest that patients classified as high-risk by InvGraphCox have a significantly greater likelihood of experiencing adverse events compared to those classified as low-risk. This highlights the model’s effectiveness in identifying patients who may benefit from more aggressive therapeutic interventions. Overall, these results underscore the potential of InvGraphCox as a valuable tool for enhancing prognostic accuracy and informing clinical decision-making in breast cancer management.

Additionally, to validate the biological interpretability of our model, we compared the top 10 clinical variables ranked by coefficient significance (Figure 7G). In breast cancer clinical practice, ER (Estrogen Receptor), PR (Progesterone Receptor), and HER2 are the canonical biomarkers that define molecular subtypes and guide therapy (13). Strikingly, InvGraphCox successfully recovered all three key receptors (ER, PR, HER2) among its top 10 most significant features. In contrast, Stable Cox identified only ER status. This indicates that, despite the challenges of mixed-type clinical data, the graph-based regularization of InvGraphCox aligns more closely with established oncological biology, prioritizing variables that are known to drive disease progression.

## Discussion

In this study, we introduced InvGraphCox, a framework designed to bridge the gap between high-dimensional biomedical data and robust survival prediction in multi-center settings. While the Cox PH model and its deep learning variants have become standard tools in oncology, their reliance on the i.i.d. assumption renders them vulnerable to distribution shifts common in heterogeneous patient cohorts. By integrating graph-structured representation learning with cohort-wise invariant alignment, InvGraphCox moves beyond treating patients as isolated instances. Instead, it leverages the intrinsic topological structure of patient populations to extract prognostic signals that are biologically stable and generalizable across different patient populations.

Our extensive benchmarking on transcriptomic data (HCC, Breast, Melanoma, and Ovarian cancer) demonstrates that InvGraphCox consistently outperforms state-of-the-art baselines, including Stable Cox and standard Cox PH models. The superior performance in curated top-100 and high-dimensional settings suggests that the patient-patient similarity graph captures latent biological manifolds that linear reweighting methods (like Stable Cox) may miss. By employing a VGAE with GraphSAGE convolution, our model aggregates information from a patient’s “genomic neighbors,” effectively smoothing out technical noise and individual outliers. This is particularly evident in the ablation study, where the removal of the GNN component resulted in a marked performance drop, confirming that the topological context is as critical as the feature values themselves for accurate risk stratification.

A central challenge in multi-center studies is preventing models from learning “site-specific” shortcuts, which are features that correlate with survival in one hospital due to batch effects but fail in another. Our invariance diagnostics (Site-AUC, HSIC, and Silhouette scores) confirm that InvGraph-Cox successfully strips away cohort-identifiable information from the latent embeddings. Crucially, this statistical invariance translates into biological stability. The high Jaccard overlap of top-ranked genes across disjoint test cohorts indicates that the model is identifying a core set of universal prognostic biomarkers rather than overfitting to training-specific idiosyncrasies. Furthermore, the gene-to-risk heatmap reveals that InvGraphCox maintains consistent directionality (protective vs. hazardous) for key genes across cohorts, a property essential for clinical trust and reproducibility.

An important observation from our results is the performance disparity between genomic and clinical data. While InvGraphCox achieved substantial gains in transcriptomic datasets, its advantage over Stable Cox was narrower in the clinical datasets (Lung and Breast cancer). We attribute this to the nature of the input space. Transcriptomic data is continuous and high-dimensional, allowing for the construction of dense, meaningful Euclidean neighborhoods that reflect biological similarity. In contrast, clinical data often consists of mixed-type covariates with sparse one-hot encodings (e.g., disease stage, treatment codes). In such spaces, Euclidean distance may fail to capture true prognostic similarity, leading to a graph structure that reflects shared administrative codes rather than shared survival trajectories. This suggests that while graph propagation is powerful for dense molecular data, its utility saturates when applied to low-dimensional, sparse clinical variables where the “manifold assumption” is weaker.

Despite its robustness, InvGraphCox has limitations. First, the graph construction step relies on a fixed *k* and Gaussian kernel bandwidth, which may not be optimal for all dataset sizes or densities. Adaptive graph learning, where the adjacency matrix is optimized jointly with the encoder, could address this. Second, as noted, the current distance metric is suboptimal for mixed-type clinical data; future iterations could incorporate learnable metrics or heterogeneous graph structures to better handle tabular data. Finally, while cohort-wise residualization is effective for mean shifts, it assumes that the underlying covariance structure of the data remains relatively stable.

In conclusion, InvGraphCox represents a significant step forward in robust survival analysis. Formalizing the patient population as a graph and enforcing distributional invariance offers a reliable alternative to standard methods that struggle with heterogeneity. As biomedical research increasingly relies on large-scale, multi-site consortia, frameworks like InvGraphCox will be essential for translating high-dimensional data into generalizable clinical insights.

## Methods

### Algorithm of InvGraphCox

#### Problem Formation

We present InvGraphCox, a graph-based survival modelling framework that integrates patient-patient relational structure with invariant representation learning to improve robustness under distribution shifts across cohorts. The model consists of three sequential components. It begins by constructing a patient-patient graph from transcriptomic profiles, where k-nearest-neighbor relationships combined with a Gaussian kernel define the weighted adjacency matrix. Next, a Variational Graph Autoencoder (VGAE) is trained to learn low-dimensional and graph-aware patient embeddings. Before survival modelling, each test cohort undergoes a cohort-wise residualization procedure that aligns its embedding mean to the training distribution to reduce inter-cohort distributional shifts. Then, a Cox proportional hazards model is fitted on the training embeddings, and its learned coefficients are applied to the aligned test embeddings to produce risk scores and survival predictions. Finally, we map the genes in the latent embedding back to their risk scores for the correlation analysis.

Suppose we have a patient-by-gene transcriptomic matrix ***X*** ∈ ℝ^*N* ×*M*^, where *N* denotes the number of patients and *M* the number of genes. For each patient *i*, the survival outcome is given by the observed time *t*_*i*_, and an event indicator *δ*_*i*_ ∈ {0, 1}, where 1 means an observed event and 0 indicates censoring. For each transcriptomic survival dataset, we first standardize the survival outcomes and gene matrix. Overall survival is defined using (*t*_*i*_, *δ*_*i*_), which are treated as outcome metadata, whereas all remaining columns of ***X*** correspond to gene expression features. The details of the three stages of InvGraphCox’s framework are in the following:

#### Graph Construction and Representation Learning

We represent each cohort as a patient-patient k nearest neighbor graph constructed from the gene expression matrix ***X*** ∈ ℝ^*N* ×*M*^. For each cohort, we compute pairwise Euclidean distances between patients’ expression vectors. This yields a directed KNN graph. We then convert KNN distances into continuous similarity weights using a radial basis function (RBF) kernel 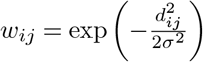, so that more similar patients receive larger edge weights. The adjacency matrix is then symmetrized to obtain an undirected weighted graph **A** = [*w*_*ij*_]. After that, we compute the degree matrix **D** = diag(*d*_*i*_) with *d*_*i*_ = ∑_*j*_ *w*_*ij*_, and obtain a symmetrically normalized adjacency

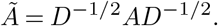

This normalization stabilizes the scale of neighborhood aggregation across nodes and is the matrix used by the graph encoder later.

To learn low-dimensional patient representations that capture the graph structure, we train a Variational Graph Auto Encoder (VGAE) (20) with a GraphSAGE encoder. The encoder consists of two GraphSAGE convolutional layers followed by two separate linear heads that output the posterior mean *μ* ∈ ℝ^*N* ×*d*^ and log-variance log *σ*^2^ ∈ ℝ^*N* ×*d*^ for each node’s latent embedding. The VGAE uses an inner-product decoder to reconstruct adjacency:

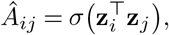

where *z*_*i*_ is sampled via the reparameterization trick *z*_*i*_ = *μ*_*i*_ + *σ*_*i*_ ⊙ *ε*_*i*_, *ε*_*i*_ ∼ *N* (0,*I*). We take the posterior means *μ* as final embeddings *Z* ∈ ℝ^*N* ×*d*^.

#### Invariant Embedding and Survival Modeling

To mitigate cohort-specific mean shifts in the learned embeddings, we apply an alignment before fitting the survival model. Let 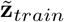 denote the mean embedding vector over the training cohort and 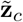 the mean embedding for a given test cohort *c*. For each patient *i* in *c*, we define a recentered embedding

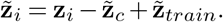

This operation removes cohort-specific mean shifts while preserving within-cohort variation, and is applied separately to each test cohort.

We then fit a penalized Cox PH model on the training embeddings. For a patient with embedding vector *x* ∈ ℝ^*d*^, the Cox model specifies the hazard:

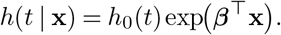

Here, we apply two-stage fitting with feature selection. We first fit the Cox model on all *d* embedding dimensions of the training cohort to obtain coefficient estimates and corresponding Wald p-values for each dimension. Then, We rank embedding dimensions by their *p*-values and select the top-k most prognostic dimensions (smallest p-values). Finally, we refit a Cox model using only these top-k features and use this reduced model for prediction on the test cohorts.

#### Biomarker Risk Attribution and Interpretation

After training the VGAE encoder, each patient’s raw expression vector *x* ∈ ℝ^*N* ×*M*^ is mapped into a latent embedding *z* ∈ ℝ^*d*^. This encoder is nonlinear and graph-aware, meaning it uses both gene expression levels and the graph structure to create embeddings *z* that summarize important biological and survival-related patterns. However, the encoder is a black box, and this means that we cannot directly interpret how each gene contributes to each latent dimension.

To analyze it, we apply a linear probe:

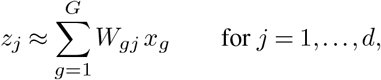

where *x*_*g*_ is expression level of gene *g, W*_*gj*_ is learned regression coefficient of *g* for latent dimension *z*_*j*_. We fit this mapping using cross-validated ridge regression (L2 regularization) to ensure stability. The result is a gene to latent mapping matrix *W* ∈ ℝ^*G*×*d*^.

We then map the latent *z* (linear probe) to risk *r*. The Cox proportional hazards head predicts the risk score 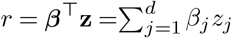. Now, we can compose the linear probe (coefficients *W*_*gj*_) with the Cox head *r* to get gene-level risk weights:

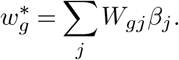

Here, if 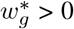: gene is risk increasing (Hazardous). If 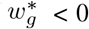: gene is risk decreasing (protective), otherwise, gene is null effect.

### Experiment Settings

#### The Top-100 Gene Benchmark Data

Using StableCox-preprocessed data (top-100 genes and OS) (12), we further evaluated cross-cohort performance across three cancer types. The HCC data (5 cohorts) used TCGA-LIHC (21) (*n* = 351) for training and Grinchuk et al. (22) (*n* = 115) for validation, generalization testing on Fujimoto (23), Roessler (24), and Hoshida (25) (*n* = 203, 209, 80). The Breast Cancer data (5 cohorts) designated 763 and 170 patients for training and validation (26), respectively, with three disjoint Curtis et al. cohorts) (26) (*n* = 521, 288, 238) serving as test sets. The Melanoma datasets (5 cohorts) was trained on Liu et al. (27)(*n* = 120) and validated on Hugo et al. (28) (*n* = 26), with testing on the Gide (29), Riaz (30), and Van (31) cohorts (*n* = 91, 54, 41).

For the univariate Cox regression analyses on the clinical subgroups for the Breast Cancer data, we follow the same setting in StableCox. We fitted an univariate Cox model to compare the high-risk and low-risk groups among patients, with the hazard ratio (HR) and 95% confidence interval (CI). Furthermore, we use a univariate Cox regression model with the selected gene as the sole covariate and computed the HR value and a log-rank test *p*-value. We label the result per cohort for a gene as favorable if HR < 1 and P value < 0.05, unfavorable if HR > 1 and P value < 0.05, and no trend otherwise. For each gene, we then calculate the percentage of cohorts falling into each category. If the genes show both favourable and unfavourable relationships with the survival outcome, it underscores that the genes screened by the model exhibit unstable prognostic relationships at the individual gene level.

#### The High-Dimensional Transcriptomics Benchmark Data

To evaluate survival models under cross-cohort distribution shifts, we utilized data from ovarian cancer (OV) and breast cancer (BRCA).

The OV compilation (32) contains 22 datasets with overall survival (OS) information, which Gao et al. grouped into 10 cohorts based on sequencing platform: GPL96 (*n* = 409)(33–36), GPL570 (*n* = 597)(37–42), GPL2986 (*n* = 194)(43), GPL6480 (*n* = 676)(44–47), GPL7264 (*n* = 128)(48), GPL7759 (*n* = 157)(49), GPL8300 (*n* = 42)(38), GPL16791 (*n* = 84)(50), OV-AU (*n* = 93)(51), and TCGA-OV (*n* = 562)(21). We utilized TCGA-OV as the training set and assessed generalization performance on the remaining nine independent test cohorts.

The BRCA data (*n* = 1, 443) spans five cohorts with gene expression and OS outcomes. We designated Yau et al.’s dataset (*n* = 682) for training (19), Caldas et al.’s dateset (*n* = 113) for validation (52), and the remaining three cohorts (*n* = 236, 295, 117) for external testing (53–55).

To facilitate cross-cohort analysis, we harmonized the independent transcriptome cohorts for each cancer type by intersecting their gene sets, restricting all data to a common feature space with *M* genes. We addressed missing expression entries *x*_*i,g*_ in the aligned data via gene-wise mean imputation performed independently within each cohort: 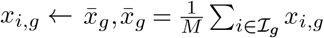, where ℐ_*g*_ denotes the indices of observed samples for the given gene.

#### The Clinical Survival Benchmark data

We evaluated the model’s generalizability on publicly available clinical cohorts for lung cancer and breast cancer (data sourced from Stable Cox) (12), aiming to demonstrate consistent prognostic performance across diverse patient subgroups and independent sites. For the lung cancer cohort, the data split was maintained identically to the Stable Cox baseline: the training set comprised 40% of patients (*n* = 240). The remaining patients were divided into validation and testing groups based on clinical criteria. The validation set was defined by sex subgroups (Female, *n* = 4; Male, *n* = 152). The testing sets were composed of non-overlapping clinical subgroups based on: age (≤60, *n* = 43; *>* 60, *n* = 113), tumor location (Central, *n* = 47; Peripheral, *n* = 107), and presence of obstructive pneumonitis/atelectasis (Present, *n* = 80; Absent, *n* = 76). The survival endpoints were Overall Survival (OS) and Disease-Free Survival (DFS). For the breast cancer cohort, the dataset was partitioned into one training cohort (*n* = 394), one validation cohort (*n* = 14), and two independent testing cohorts (*n* = 273 and *n* = 177). The survival endpoints were OS and RFS.

Unlike the all-numeric, pre-scaled features used for the transcriptome data, the clinical cohorts contain mixed-type covariates (including continuous, binary, and one-hot encoded features). This necessitated an adapted feature preprocessing and distance calculation procedure for the *k*-Nearest Neighbor (KNN) graph. We strictly adhered to the “train-once, test out-of-distribution” protocol to prevent data leakage and ensure robustness against cohort shifts. A single `ColumnTransformer` was exclusively fit on the training cohort to perform the following steps: (i) Impute missing values (mean for continuous, most frequent for categorical/binary), (ii) Z-score scale continuous features to ensure the Euclidean distance is meaningful, and (iii) remove features with zero variance. This frozen transformer was then applied, without refitting, to all validation and test cohorts. Following feature standardization, we constructed a patient KNN graph in the standardized clinical feature space using Euclidean distance. The distance matrix *d*_*ij*_ was converted into RBF edge weights 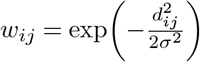. The bandwidth parameter *σ* was determined solely from the KNN distance distribution of the training set. Crucially, this *σ* value was then reused unchanged to construct the graphs for all validation and test cohorts, thereby ensuring comparable edge scales across all datasets.

Afterwards, the same graph representation learning approach (VGAE+GNN) used in the transcriptome experiment was applied to the constructed clinical graphs. A Cox Proportional Hazards (Cox PH) model was subsequently trained exclusively on the latent embeddings derived from the training cohort. For validation and testing, the trained Cox PH model produced log-partial hazards from the respective test embeddings. We quantified the prognostic performance by computing the C-index for each independent test set and reported the overall mean C-index across all testing cohorts.

To compare the clinical variables prioritized by InvGraphCox and Stable Cox (as presented in the radar plot), we adopted the coefficient significance ranking methodology from Stable Cox. The *p*-value (*P*) of each variable’s coefficient was transformed into a measure of statistical significance using the transformation −log_2_(*P*). These transformed, scaled values were then used as the radial coordinates of the radar chart, enabling a visual comparison of the relative prognostic relevance of clinical features between the models.

#### Baseline Methods

We benchmark InvGraphCox against five established survival models: the Stable Cox model (a distribution-shift-robust variant of Cox) (12) and the Cox Proportional Hazards (Cox PH) model (3), along with three parametric Accelerated Failure Time (AFT) models: Weibull (7), log-logistic (8), and log-normal (9) models. This comparison evaluates our model’s performance and robustness in comparison to standard and state-of-the-art methods for prognostic prediction.

##### Stable Cox

Stable Cox addresses covariate shift by first learning sample weights that decorrelate covariates, then fitting a weighted Cox PH on the reweighted empirical distribution. First is the independence-driven sample reweighting. Let 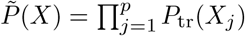 be the product of marginal (re-sampled) feature distributions, where *P*_tr_ is the training distribution. The model learn the density-ratio weights:

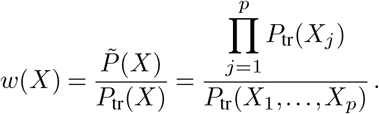

Intuitively, this enforces independence among covariates in the weighted sample. Second is the weighted Cox partial log-likelihood. With weights *w*(*x*^(*i*)^), the (negative) loss uses the weighted partial log-likelihood:

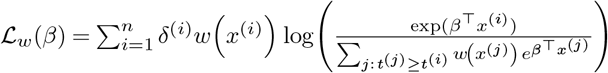

Under mild conditions, coefficients on unstable variables shrink to 0. We implement Stable Cox as described in the original paper’s setting (12), which involves a two-stage framework: SRDO weighting and weighted Cox.

AFT models describe survival times via time scaling:

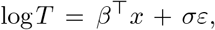

so covariates accelerate time by factor exp(*β*^⊤^*x*). Below, *S*_0_, *h*_0_ denote baseline survival function and hazard.

##### Weibull AFT

In Weibull AFT, the hazard is monotone (decreasing for *k* < 1, constant for *k* = 1, increasing for *k* > 1), making it good when risk changes in one direction over time. With shape *k* > 0 and scale *λ* > 0:

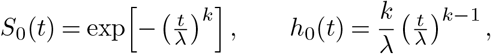

where *k* controls the trend of the hazard, it is the shape of how risk changes over time, and *λ* is the time scale.

##### Log-logistic AFT

In log-logistic AFT, the hazard is typically non-monotone, meaning that risk can rise and then fall, and the distribution has a heavy tail, so late events remain relatively likely. With shape *k* > 0 and scale *λ* > 0:

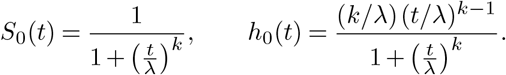

##### Log-normal AFT

In log-normal AFT, log *T* is approximately Gaussian, producing a hazard that rises and then de(clines wi th a moderately heavy tail. With log *T* ∼ *N*(*β* ^⊤^*x, σ*^2^):

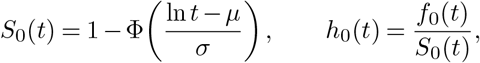

here, the *f*_0_(*t*) is the density function 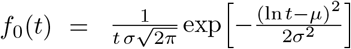.

#### Patient Risk Evaluation Metrics

The performance of the survival model’s patient risk scores was primarily quantified using two summary metrics: the Concordance Index (C-Index) and the Integrated Brier Score (IBS). The Kaplan-Meier (KM) survival curves were also used as a key visualization tool to assess risk stratification ability.

#### Concordance Index (C-Index)

The Concordance Index (C-Index) is a rank-based metric representing the probability that, for a randomly chosen comparable pair of patients, the individual with the shorter observed survival time is correctly assigned a higher predicted risk.

Let each patient *i* have observed time *t*_*i*_, event indicator *δ*_*i*_ ∈ {0, 1}, and a scalar risk score *r*_*i*_. The set of comparable pairs *P* (Harrell’s *C*) is defined as:

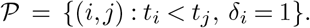

The C-index estimator is:

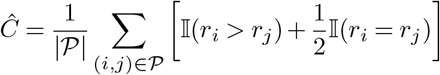

For each testing cohort, we evaluated performance by computing Harrell’s C-index, utilizing the model’s predicted partial hazards as the risk score *r*_*i*_.

#### Kaplan-Meier Survival Curves

To visually assess the model’s ability to stratify patient risk, we generate Kaplan-Meier (KM) survival curves for each external test cohort. Patients are divided into high-risk and low-risk groups based on their predicted log-partial hazard (linear predictor) from the trained Cox model. The median value from the training cohort is used as a fixed cutoff to define these strata. Patients with predicted risks above this cutoff are assigned to the high-risk group; those below are assigned to the low-risk group. We then estimate separate KM survival curves for the two strata. The statistical significance of the separation is assessed using the log-rank test. Furthermore, we quantify the relative risk by fitting a univariate Cox model using only the binary risk group indicator to estimate the Hazard Ratio (HR) and its 95% confidence interval.

#### Integrated Brier Score (IBS)

The Integrated Brier Score (IBS) serves as a robust measure of overall time-dependent prediction accuracy. At any time *t*, the time-dependent Brier score BS(*t*) is the mean squared error between the observed event-free indicator *Y*_*i*_(*t*)= **1**(*t*_*i*_ *> t*) and the predicted survival probability *Ŝ*_*i*_(*t*). To adjust for informative censoring bias, we apply Inverse Probability of Censoring Weighting (IPCW) using the censoring survival probability *Ĝ*_*i*_(*t*) (estimated via the Kaplan-Meier method on censoring times), where *w*_*i*_(*t*) are the IPCW weights.

The Brier score at time *t* is:

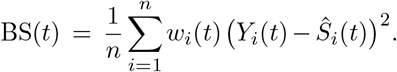

The IBS is then calculated by integrating BS(*t*) over a time window [*τ*_0_, *τ*_1_]:

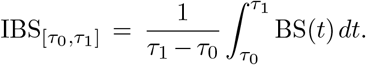

#### Choice of the Time Window

The integration window [*τ*_0_, *τ*_1_] is defined across the three testing cohorts to ensure computational stability and robustness. The bounds are set as *τ*_0_ = max(1st-percentiles of min *T*) and *τ*_1_ = min(max *T*), where *T* denotes the observed follow-up times. We select the maximum of the 1% follow-up time percentiles for *τ*_0_ because starting slightly later stabilizes the IPCW estimation, mitigating high variance often seen at early times due to a limited risk set and irregular censoring. This choice ensures that approximately 99% of each cohort remains under observation at the start time *t*_lower_.

#### Uncertainty via Bootstrap

The uncertainty in the IBS estimate is quantified by reporting the 95% confidence interval (CI) derived using a nonparametric bootstrap procedure. The cohort is resampled *B* times, and the IBS estimate 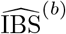 is recomputed for each replicate. The percentile CI is then constructed using the 2.5^th^ and 97.5^th^ percentiles of the ordered bootstrap replicates *θ*^*^ of 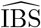:

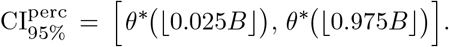

This interval reflects the precision of the IBS estimate for the specific cohort and prediction horizon.

#### Invariance Diagnostics

To demonstrate the model’s robustness to covariate distribution shift (i.e., changes in *P* (*X*) across different hospitals, platforms, or populations), we employed a set of invariance diagnostics. While the feature distribution *P* (*X*) is expected to change in cross-cohort scenarios, the underlying biological mechanism, represented by the conditional distribution *P* (*Y* | *X*), should remain stable. A prognostic model is truly generalizable only if it learns features that are invariant to cohort-specific noise. We test two complementary properties across all test cohorts:

#### Cohort Invariance (No Site Leakage)

This property ensures that the model’s outputs, both the final risk scores *r* = *β*^⊤^*z* and the intermediate latent embeddings *z* ∈ ℝ^*d*^, are statistically independent of the source cohort identity. This confirms that the model avoids learning spurious, cohort-specific correlations. The metrics used to quantify this invariance are Site-AUC(*r*) and Site-AUC(*z*), which quantify the model’s ability to classify the source cohort using the risk score *r* or the embedding *z*. A value close to 0.5 is desired. Hilbert-Schmidt Independence Criterion (HSIC(*z*, site))(16): Measures the statistical dependence between the latent embedding *z* and the cohort label. A value near zero is ideal. Silhouette Score (Silhouette(*z* | site))(17): Assesses clustering quality in the latent space; a value near zero or negative is favorable, indicating patients from different cohorts are mixed well.

#### Biological Stability (Shared Prognostic Signal)

This property verifies that the core biological mechanisms learned are consistent across cohorts, despite shifts in covariates. Specifically, we require the learned gene-to-risk associations to remain stable. The consistency is evaluated using: Top-*K* Jaccard overlap of important genes. Genome-wide rank Spearman correlation of prognostic scores. Top-*K* signed Spearman heatmap (meta-genes) for visual confirmation.

In summary, the Cohort Invariance diagnostics validate the removal of spurious site signals, ensuring that risk and embeddings do not carry cohort-specific information. Concurrently, the Biological Stability analysis confirms that the model successfully maintains the fundamental, generalizable prognostic mechanisms that drive generalization.

## Data Availability

The platform-level external cohort ovarian cancer data were acquired from Gao et al.’s study (32), and the breast cancer data were fetched from the Xena platform (56). Both the top-100 gene benchmark and the clinical benchmark data were retrieved from Stable Cox (12).

## Code Availability

The code is available at https://github.com/compbioclub/InvGraphCox.

## Acknowledgments

We express our gratitude for the support provided by the National Natural Science Foundation of China (No. 32400519), the Research Grants Council of Hong Kong (No. 21200425), the CityUHK Start-Up Grant (No. 9610687), and the Tung Biomedical Sciences Centre Project Fund (No. 9609331). We acknowledge the use of ChatGPT-5 and Gemini-3 for grammatical and stylistic refinement of the text. We acknowledge that specific graphical elements within Figure 1 were generated using the AI tool Nano Banana Pro, based on prompts designed by the authors. No confidential or sensitive data was shared during this process. All AI-assisted outputs were critically reviewed and approved by the authors, who retain full responsibility for the content.

## Author Contribution Statements

KHN: Software; Formal analysis; Validation; Visualization; Investigation; Methodology; Writing—original draft. CSL: Formal analysis; Validation; Visualization, Writing—original draft. ANJ: Validation; Investigation; Writing—review & editing. YHL: Investigation; Writing—review & editing. LXC: Conceptualization; Supervision; Formal analysis; Visualization; Methodology; Writing—original draft.

